# Dual Knockout of *StAMY23* and *StVINV* Improves Postharvest Storage Traits in Potato

**DOI:** 10.64898/2026.06.08.730856

**Authors:** Paula Teper-Bamnolker, Talya Steinberg, Carmela Shtein, Reut Peer, Adi Doron-Faigenboim, Eduard Belausov, Amir Sherman, Dani Eshel

## Abstract

Starch is the primary carbohydrate reserve in potato (*Solanum tuberosum* L.) tubers and a critical determinant of their industrial value. The rate of starch degradation during postharvest storage directly influences key traits such as endodormancy (ED) duration and cold-induced sweetening (CIS), which affect sprouting behavior. In this study, we used CRISPR/Cas9 genome editing to knockout *StAMY23*, a gene encoding α-amylase involved in starch breakdown. *stamy23* plants exhibited higher yield and extended tuber ED postharvest, without significantly altering CIS or starch granule content. To further reduce CIS, we knockout *StAMY23* in *VACUOLAR INVERTASE* knockout *(stvinv*) backgrounds, generating *stamy23/stvinv* double-knockouts plants. These lines showed significantly reduced CIS, prolonged ED, and elevated starch content, along with altered starch granule content. Collectively, our findings demonstrate that simultaneous downregulation of *StAMY23* and *StVINV* can additively enhance desirable postharvest traits, providing a promising strategy for improving potato storage quality through precision genome editing.

## Introduction

Potato (*Solanum tuberosum* L.) is the fourth most important agricultural crop globally, following rice, wheat, and maize. It is consumed both fresh and in processed forms, generating consistent, year-round demand that necessitates long-term post-harvest tuber storage [1, 2]. The nutritional value of potato tubers contributes to their global significance; they are an excellent source of carbohydrates and contain essential minerals, vitamins, and amino acids [3]. In addition to its role as a staple food crop, the potato is extensively cultivated for starch production. Potato-derived starch is widely utilized across the food and industrial sectors and is frequently modified chemically or physically to meet specific functional requirements [4]. The dry matter and starch contents are critical determinants of tuber quality, particularly with respect to their suitability for consumption, processing, and industrial starch extraction [5].

Potato tubers contain approximately 13.5–15% starch on a fresh weight basis and 75–80% starch on a dry weight basis [6]. Starch biosynthesis occurs during tuber development; Sucrose, transported from photosynthetically active leaves via the phloem, is converted into starch within the tuber’s amyloplasts, organelles that deposit starch as a water-insoluble mixed-glucose polymer. This polymer consists of amylose (∼25%), a primarily linear polymer of α-1,4-linked glucose units, and amylopectin, a branched polymer composed of α-1,4-linked glucose chains with α-1,6-linked branch points [7]. Functionally, starch serves dual roles in plant metabolism: it acts both as a carbon reserve that is mobilized when energy demands exceed supply and as a storage depot when carbon assimilation exceeds short-term requirements, thereby enabling efficient carbon utilization [3].

Harvested tubers possess buds that remain dormant and will not sprout, even under optimal sprouting conditions [8, 9]. This physiological state, known as endodormancy (ED), is regulated by unidentified endogenous signals that suppress meristem growth [10, 11]. The ED period is influenced by genotype, crop growth conditions, and postharvest storage, correlating more closely with “sugar unit” accumulation than with chilling units [12]. After a transition period of 1 to 15 weeks, dormancy is broken, and the apical bud resumes growth [13, 14]. To delay sprouting and reduce losses due to diseases, postharvest tubers are typically cold stored. This ensures a stable year-round supply for consumers and the food processing industry [15]. However, cold storage induces a process known as cold-induced sweetening (CIS), a phenomenon marked by the accumulation of reducing sugars (glucose and fructose) as a result of starch degradation. These sugars react with free amino acids during heat processing via the Maillard reaction, forming undesirable dark-colored, bitter products and the potentially harmful compound acrylamide [16, 17]. Therefore, CIS poses a significant challenge to the potato industry and raises worldwide food safety concerns [18, 19]. During cold storage that leads to CIS and the ED-to-growth transition, starch is gradually broken down into soluble sugars to meet the energy and metabolic demands of sprouting and stress adaptation. This degradation is controlled by a complex regulatory network involving multiple enzymes and signaling pathways. Among these, alpha-amylase 23 (StAMY23) and vacuolar invertase (StVINV) enzymes are key regulators [20-22].

Alpha-amylases are endoamylolytic enzymes that specifically hydrolyzes α-1,4-glucan bonds to form various linear and branched malto-oligosaccharides from starch, and StVINV catalyzes the hydrolysis of sucrose into glucose and fructose within the vacuole and is considered a major contributor of CIS [23, 24]. Five *StAMY* genes have been identified in the potato genome [3]. Two of them, *StAMY1* and *StAMY23*, are expressed in tubers, but only StAMY23 enzyme is induced by low temperatures [3, 20]. StAMY23 is localized in the cytoplasm and regulates CIS of tubers through the degradation of cytosolic phytoglycogen. Silencing *StAMY23* yielded a higher accumulation of phytoglycogen and lower hexose in cold-stored tubers, while the starch content was not obviously affected, suggesting that StAMY23 might operate on the soluble phytoglycogen, which is a small fraction of potato starch and is mainly deposited in the cytoplasm [20, 21]. In addition, *StAMY23*-RNAi silencing lines showed delayed tuber sprouting in storage accompanied by a higher soluble malto-oligosaccharides and decrease in the amount of glucose and fructose, as well as a slight change in the phytoglycogen structure and starch granule size [21].

Our previous work using *StVINV* CRISPR knockout and overexpression lines (35S::StVINV) demonstrated that modulating StVINV activity affects sugar unit accumulation and, consequently, the duration of tuber dormancy. *stvinv* tubers showed reduced hexose accumulation during cold storage and prolongs ED, while 35S::StVINV tubers showed the opposite response [12, 25]. In addition to CIS resistance, *stvinv* tubers exhibited significantly lower lipid peroxidation and H_2_O_2_ levels in response to cold stress, upregulating antioxidant defenses and enhancing the accumulation of raffinose family oligosaccharides. Furthermore, under cold stress and drought conditions, *stvinv* plants retained normal vigor, in contrast to wild-type plants that showed wilting symptoms [25, 26]. These findings suggest that *stvinv* plants possess enhanced tolerance to abiotic stress, presenting opportunities for genetic improvement of potato traits.

Improving starch content, reducing CIS, and extending ED duration are major objectives in potato postharvest management. In this study, we developed a genetically improved *S. tuberosum* cv. Désirée by introducing a CRISPR/Cas9-mediated knockout of *StAMY23* in a *stvinv* background. The resulting double mutants maintained improved postharvest traits compared with WT, including prolonged ED and reduced CIS, similar to the corresponding single mutants. However, the starch-excess phenotype was unique to the double mutants. This genetic combination offers a promising strategy to address major challenges in potato production by enhancing abiotic stress tolerance, increasing yield stability, and contributing to global food security.

## Results

### *stamy23* knockout induces prolonged ED and larger tubers

We hypothesize that reducing the rate of starch degradation can induce longer ED duration of potato tubers. Since *StAMY23* has previously been identified as a key gene involved in tuber starch degradation [20, 21], we aimed to perform targeted mutagenesis of all four alleles of *StAMY23* using the CRISPR/Cas9 system. To design single guide RNAs (sgRNAs) for *StAMY23* (*PGSC0003DMG400009891*), the gene sequence was analyzed using the reference potato genome (*S. tuberosum* v4.03) [27]. Two sgRNAs were selected to induce mutations in the coding regions of *StAMY23*. sgRNA1 was designed to target exon 3, a conserved region among the predicted alternative transcripts, which contains a unique *MspI* restriction site. sgRNA2 was designed to target exon 4, providing an additional site for mutagenesis and ensuring a higher likelihood of functional disruption (Fig. 1A). Following Agrobacterium-mediated transformation of Désirée cultivar, two *StAMY23* mutant lines #11 and #13 were identified through PCR analysis of the target region based on band size variations relative to WT, suggesting DNA deletions (Fig. 1B). Sequencing of the amplified products revealed various mutation types, including nucleotide substitutions, insertions, and deletions generating two *stamy23* full knock out mutant lines (Fig. 1C). Phenotypic analysis showed no variations in plant growth rate or leaf shape as compared to WT (Fig. 2A-B). However, the *stamy23#11* and *stamy23#13* displayed enhanced yield traits, characterized by the development of larger tubers (Fig. 2A-B). Dormancy analysis revealed that WT tubers reached dormancy release of 50% of the tubers after 84 days of storage, whereas *stamy23* tubers *#11* and *#13* sprouted after 90 days (Fig. 2C); Suggesting that *StAMY23* plays a role in dormancy maintenance. The levels of sucrose and hexoses showed no significant differences compared to WT (Fig. 2D). Starch analysis using safranin O staining and subsequent quantification of starch granules revealed that the single mutants *stamy23#11* and *#13* showed no significant changes in total granule density or size per amyloplast as compared to WT (Fig. 2E-G). Overall, these findings suggest that *stamy23* plants confer beneficial agronomic traits, including increased yield and extended dormancy, while also having no significant effect on CIS or starch content.

**Fig. 1.**
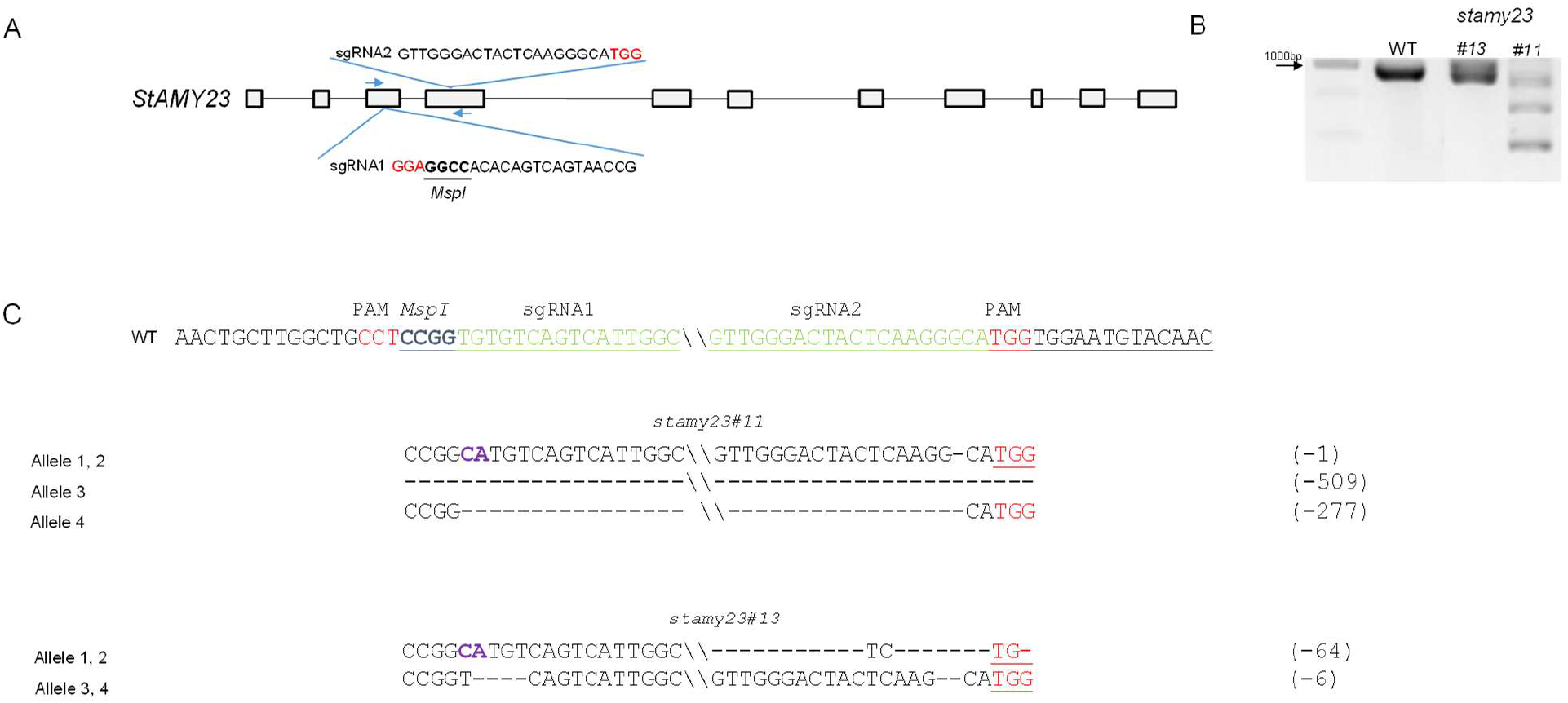
Genotyping *stamy23* CRISPR/Cas-9 lines. (A) Schematic presentation of sgRNAs target sites in *StAMY23*. The restriction site for the identification of mutated alleles is underlined, and the protospacer adjacent motif (PAM) is marked in red. Blue arrows indicate primers flanking the target region utilized for PCR amplification. (B) PCR products of Cas9-*StAMY23* target sequences from developing shoots were obtained using the primers marked in (A) by blue arrows. (C) Sequence analysis of four types of alleles. The wild-type (WT) sequence is shown at the top of each alignment, with the PAM sequences in red and the sgRNA sequence in green. Dashes indicate nucleotide deletions and substitution are marked in purple. The size of deletions (number of missing nucleotides) is specified on the right side of the sequence.

**Fig. 2.**
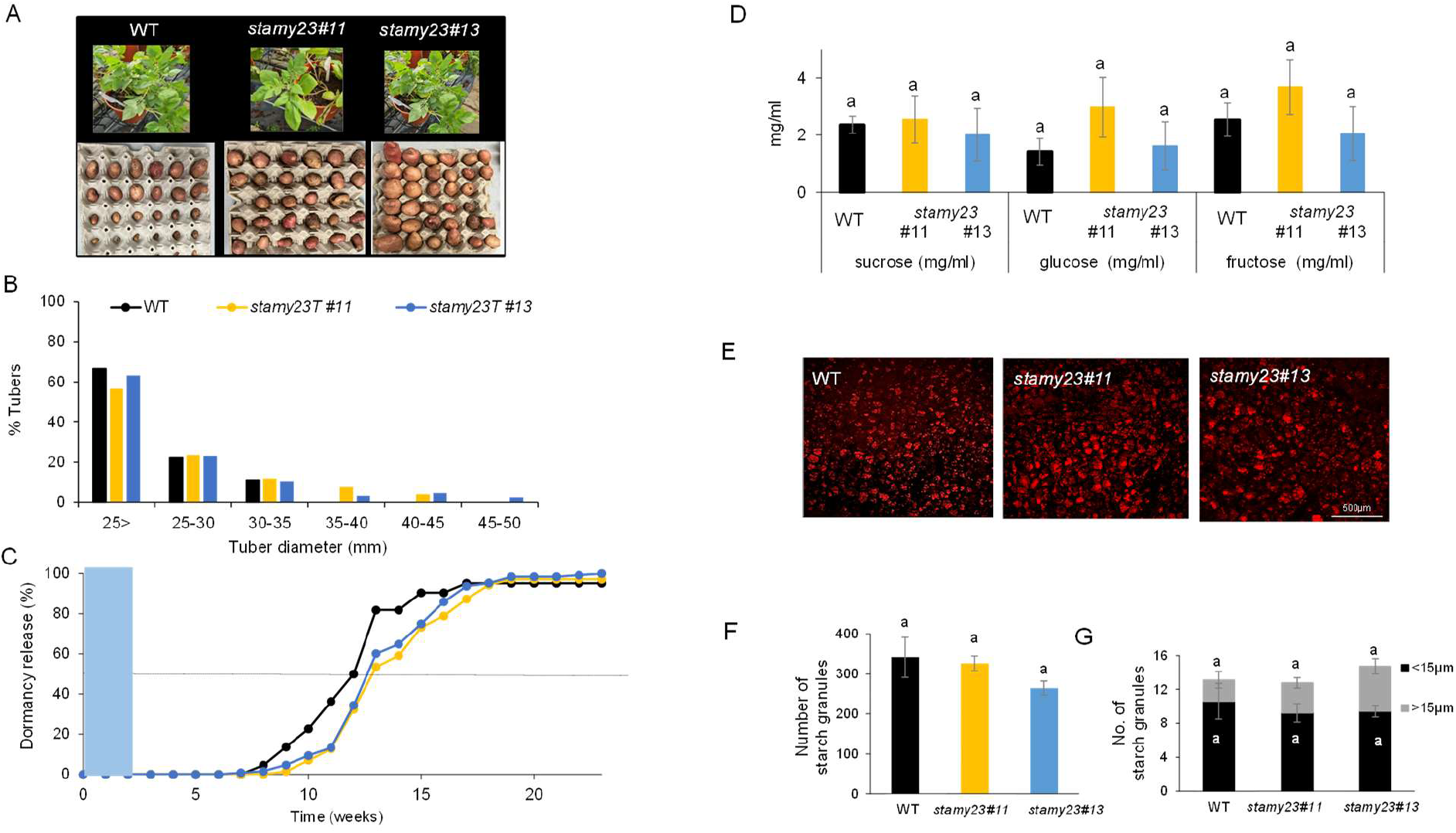
*StAMY23* CRISPR/Cas9 knockout enhances yield and prolongs tubers endodormancy without significantly affecting sugar levels or starch granule content and size. (A) Plant growth and yield of WT and *stamy23* lines. (B) Tubers size distribution. (C) The rate of endodormancy release of tubers stored for two weeks at 4°C (blue rectangle) and then transferred to 14°C, 70 tubers were analyzed for each genotype. Black dashed line indicates x_50_ time point, when dormancy of 50% of the tubers is released. (D) Sucrose, glucose, and fructose in WT and *stamy23* tubers that were stored for two weeks at 4°C followed by 10 days in 14°C. (E) Representative pictures of starch granules within amyloplasts, stained with Safranin O and observed under confocal laser microscope, 30 days after harvest, tubers were stored for two weeks at 4°C followed by 10 days in 14°C. (F) Quantification of starch granules in a square of 500µm^2^ and (G) the number of starch granules per amyloplast calculated from (E) using Imaris software. Error bar represents standard error (SE). Letters indicate significant differences between treatments at each time point (one-way ANOVA, p<0.05).

### Creation of a non-transgenic *stvinv* line for trait improvement

Our previous study demonstrated that knockout of *StVINV* via stable expression of CRISPR/Cas9 components significantly reduced CIS and prolonged ED in potato tubers [25]. We hypothesize that non-transgenic *stvinv* plants could serve as a platform for introducing additional genetic modifications aimed at further improving crop and postharvest traits. To test this, we generated *StAMY23* knockouts in the background of the non-transgenic *stvinv#21* line of cv. Désirée, developed by protoplasts transfection and regeneration using a non-binary pSAT-Cas9-sgRNA plasmid [28]. Mutant analysis using PCR amplification followed by *BsuRI* digestion revealed that the PCR product derived from *stvinv#21* was not digested (Fig. 3A). Subsequent genotyping using NGS amplicon sequencing confirmed a heterozygous plant carrying two knockout alleles and two alleles with a single–amino acid deletion (Fig. 3B, C). To evaluate the phenotype, the *stvinv#21* plants were grown under greenhouse conditions for up to 120 days. No significant differences in overall plant growth or tuber yield were observed compared to WT plants (Fig. 3D). However, after storing the tubers at 4°C, the *stvinv#21* tubers exhibited higher sucrose levels and lower hexose levels, as expected (Fig. 3E). *stvinv#21* tubers showed an extended ED period in storage, like the phenotype of the stable lines *stvinv#7* and *stvinv#8* [12]. Whereas 50% of WT tubers reached dormancy release after 90 days of storage, only 25% of *stvinv#21* tubers were dormancy releases even after 168 days (Fig. 3F). Demonstrating the successful generation of a non-transgenic *stvinv* knockout line in ‘Désirée’, which can be further utilized to produce *stamy23*/*stvinv* double mutant tubers.

**Fig. 3.**
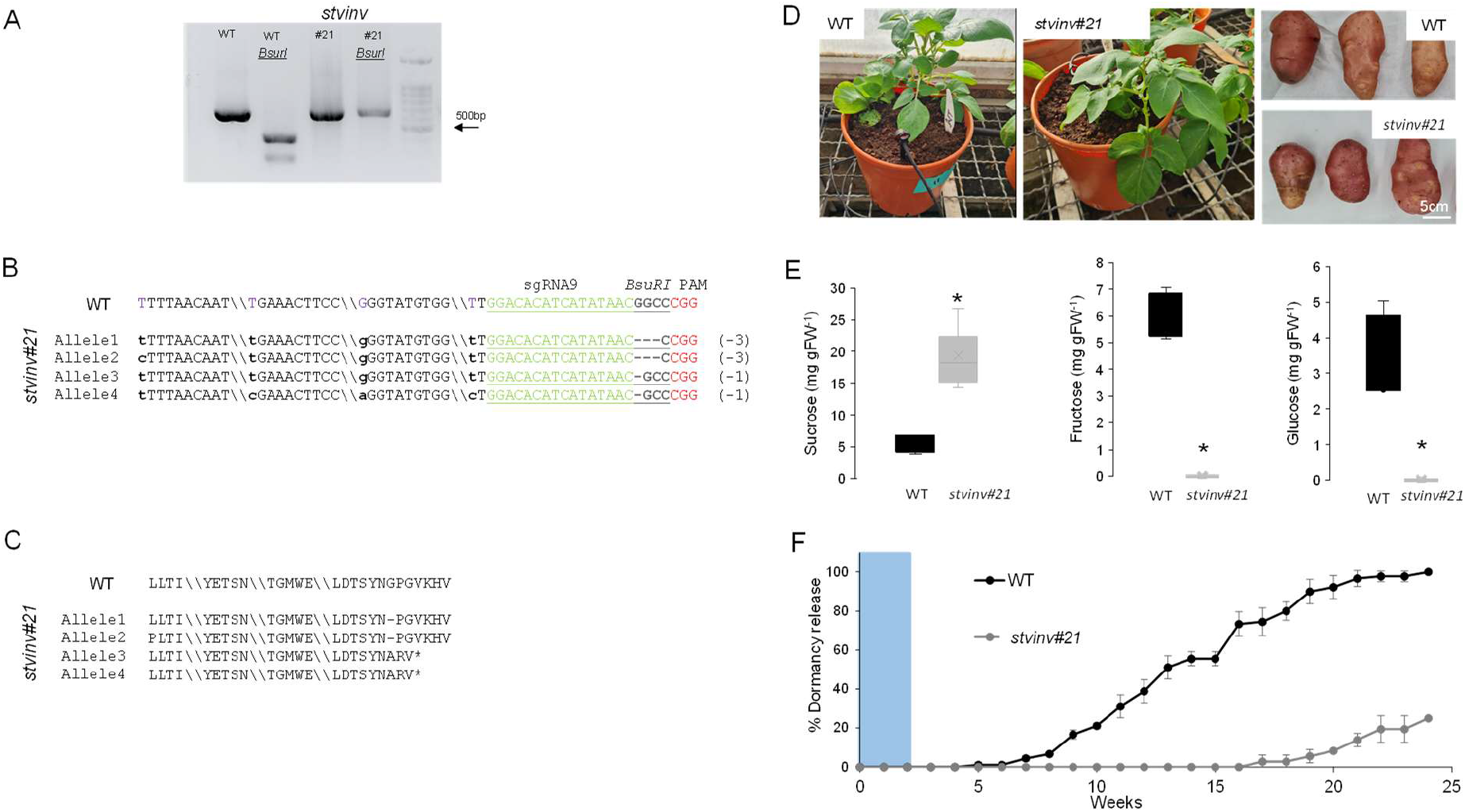
*stvinv#21* non-transgenic CRISPR/Cas9 knockout line displays reduced CIS and delays endodormancy (ED) release. (A) Restriction analysis of T0 PCR fragments of wild type (WT) ‘Désirée’ and *stvinv#21* by *BsurI* restriction enzyme. Uncut band indicates gene editing, while WT control shows a cut band. (B) Sequence analysis of four types of alleles. WT sequence is shown at the top of each alignment, with the protospacer adjacent motif (PAM; red) and sgRNA sequence (green). Deletions are represented by dashes, with the number of missing nucleotides noted on the right. Single nucleotide polymorphisms (SNPs) are shown in lowercase bold letters. (C) Predicted polypeptides from *stvinv#21* mutant line. Asterisks (*) indicate predicted stop codon, and deleted amino acids are replaced with dashes. (D) Representative WT and *stvinv#21* plants and yield following greenhouse 90 days of cultivation. (E) Sucrose, fructose, and glucose content in WT and *stvinv#21* postharvest tubers after 4 weeks at 4°C. Asterisks indicate significant differences (*P* < 0.05). (F) Percentage of dormancy release in WT and *stvinv#21* postharvest tubers stored for 2 weeks at 4°C (blue rectangle), and then transferred to 14°C. Tubers were considered dormant until at least one bud reached 2 mm length. Error bar represents standard error (SE).

### Dual knockout of *StAMY23* and *StVINV* exhibits reduced CIS, extended ED, and increased starch content

To generate double mutants, agrobacterium-mediated transformation was carried out on the *stvinv#8* and *stvinv#21* lines to knockout *StAMY23*. Following transformation, four *stamy23/stvinv#21* mutant lines (#4.1, #4.2, #5.1, and #5.3) and the *stamy23/stvinv#8* mutant line #1 were identified by PCR analysis of *StAMY23* target region (Fig. 4A). Variations in band size compared to the WT suggested DNA deletions, while the presence of an uncut band following *MspI* digestion of *StAMY23* PCR product indicated targeted mutations (Fig. 4A and B, respectively). Genotyping all the lines revealed that they contained four mutated alleles, resulting in deletions ranging from 1 to 445bp, insertions up to 272bp and the substitution of 2 nucleotides (Fig. 4C). The *stamy23/stvinv8#1* plants showed similar growth and tuber formation as compared to the WT but exhibited an extended ED in storage (Fig. 5A, B). In contrast, *stamy23/stvinv#21* lines showed growth difficulties and developed a tapered, “carrot-shaped” tubers, with a prolonged ED (Fig. 5C, D). Sugar analysis showed that *stamy23/stvinv#8* and *stamy23/stvinv#21* tubers accumulated higher sucrose and lower hexose levels following cold storage, compared to the WT tubers, resembling the sugar profile of *stvinv#8* and *stvinv-#21* (Fig. 6A, B). Starch analysis using Safranin O staining and confocal microscopy showed that the *stamy23/stvinv* double mutants displayed a starch-granule excess phenotype, defined here as increased starch-granule abundance per 500 µm^2^ area compared with WT and the corresponding *stvinv* single mutants (Fig. 7A, B; Fig. S1A–C). In the *stamy23/stvinv#8* background, this was accompanied by changes in granule-size distribution per amyloplast, including more small granules (<15 µm) and a higher proportion of larger granules (≥15 µm) relative to WT and single mutants (Fig. 7C). Chemical starch quantification in the *stamy23/stvinv#21* background showed higher starch levels in both single and double mutants compared with WT, indicating that this measurement supports increased starch accumulation relative to WT but does not by itself distinguish the double mutant from the *stvinv* single mutant (Fig. S1D).

**Fig. 4.**
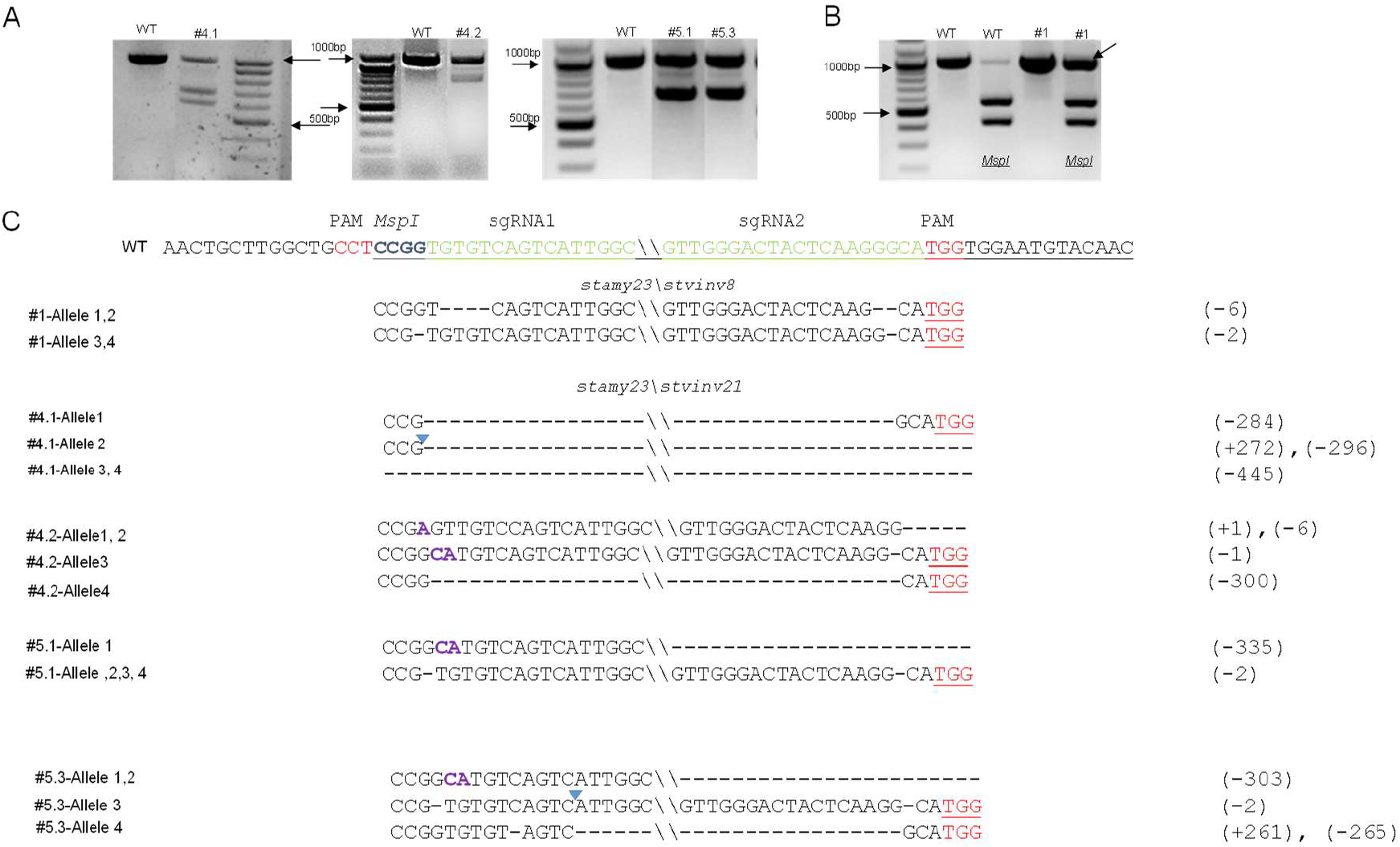
Genotyping of *stamy23/stvinv#8/#21* CRISPR/Cas9 knockout lines in Désirée cultivar. (A) PCR amplification of Cas9-*StAMY23-*sgRNA1 and sgRNA2 target sequences obtained from T0 developing shoots of *stamy23/stvinv21#4*.*1, #4*.*2, #5*.*1*, and *#5*.*3* as compared to wild-type (WT), using primers flanking the target region as indicated in Fig.1A. The presence of smaller bands relative to WT indicates DNA deletions. (B) *MSPI* restriction enzyme analysis of T0 PCR fragments, using primers flanking the target region indicated in Fig.1A. Uncut band (arrow) indicates gene editing, while WT control shows a cut band. (C) Sequence analysis of mutant alleles. The WT sequence is shown at the top of each alignment, with protospacer adjacent motif (PAM) sequences in red and sgRNA sequence in green. Nucleotide deletions are represented by dashes, insertions are marked with a blue triangle, and substitution are marked in purple. The size of deletions (number of missing nucleotides) is indicated on the right side of the sequence.

**Fig. 5.**
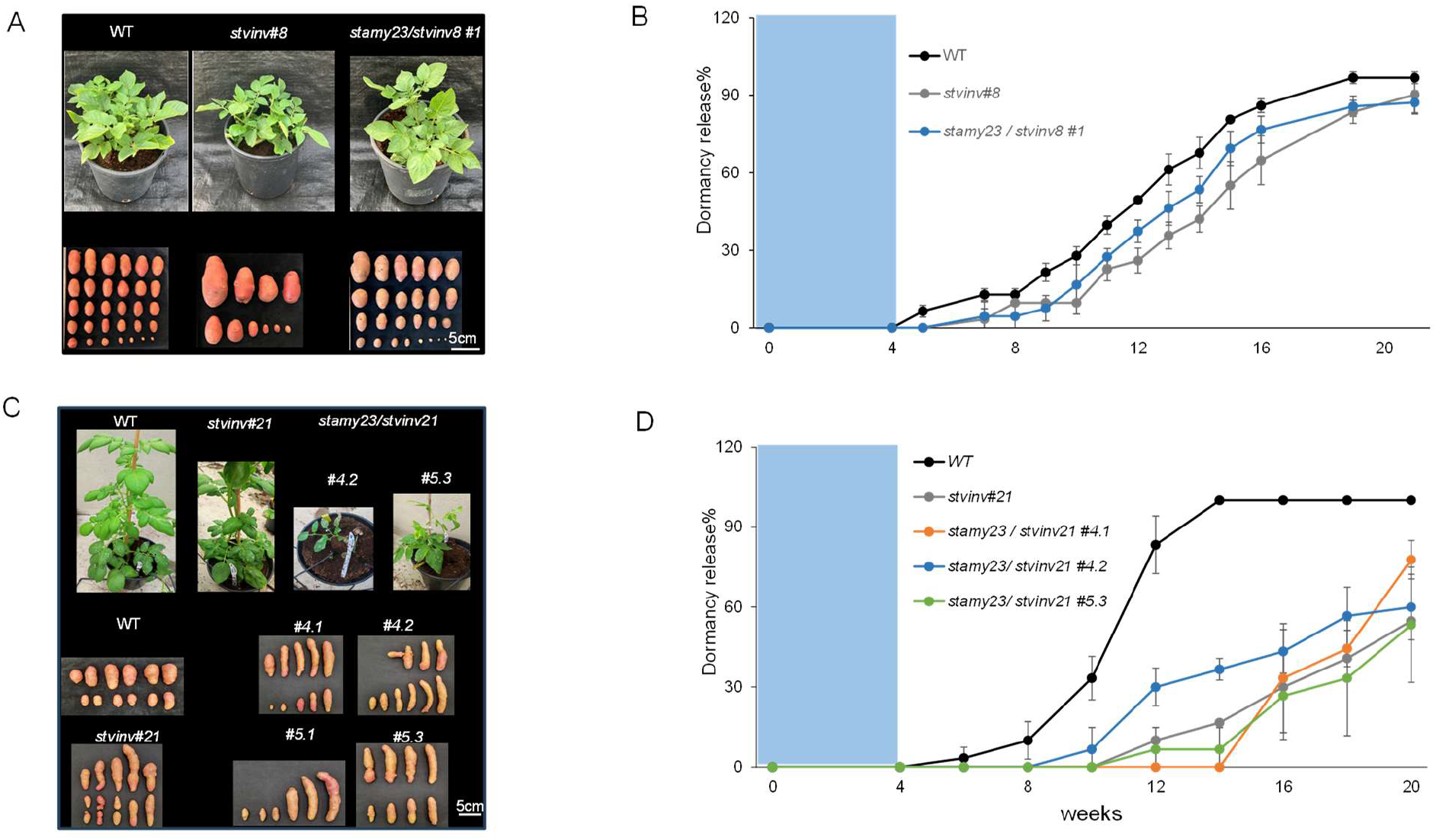
*stamy23/stvinv8* and *stamy23/stvinv21* double mutants exhibit extended endodormancy (ED) as compared to wild type (WT). (A), (C) Representative images of *stamy23/stvinv8#1* and *stamy23/stvinv21#4*.*1, #4*.*2, #5*.*1, #5*.*3* knockout lines grown in net-house for 71 or 78 days; respectively and developed tubers compared to WT and single mutants *stvinv#8/#21*. (B), (D) Tubers ED duration after 4 weeks at 4°C (blue rectangle) followed by 14°C incubation. The scale bar is equivalent to 5cm. Error bar represents standard error (SE).

**Fig. 6.**
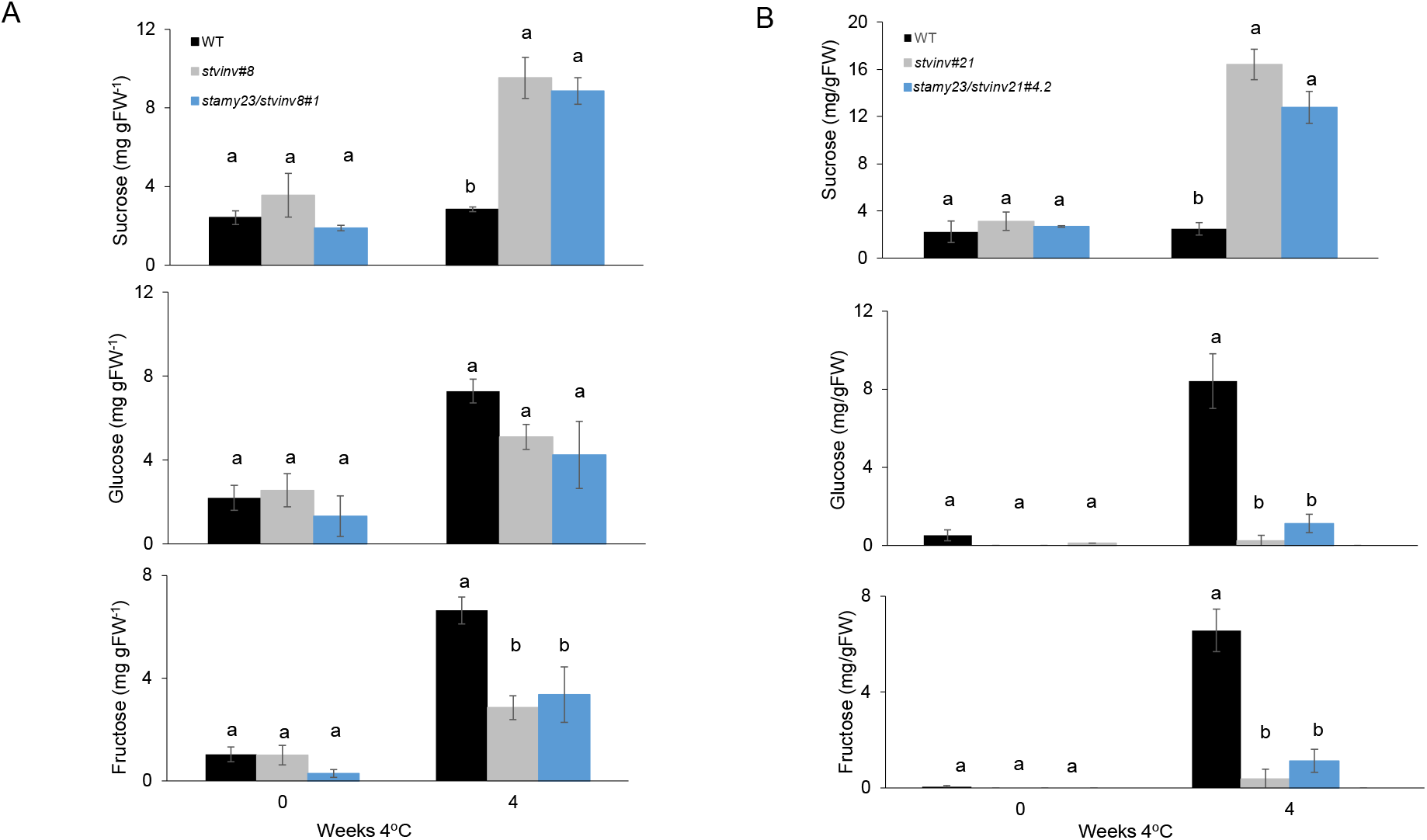
*stamy23/stvinv8* and *stamy23/stvinv21* double mutant lines exhibit reduced cold-induced sweetening (CIS). (A, B) Sucrose, glucose, and fructose levels in *stamy23/stvinv8#1*(A) and *stamy23/stvinv21#4*.*2* (B) as compared to wild type (WT) and single mutants *stvinv#8/#21* lines, during before (0) and after four weeks of exposure to 4°C. Error bars indicate standard error (SE). Different letters denote significant differences between treatments at each time point (*p* < 0.05, one-way ANOVA).

**Fig. 7.**
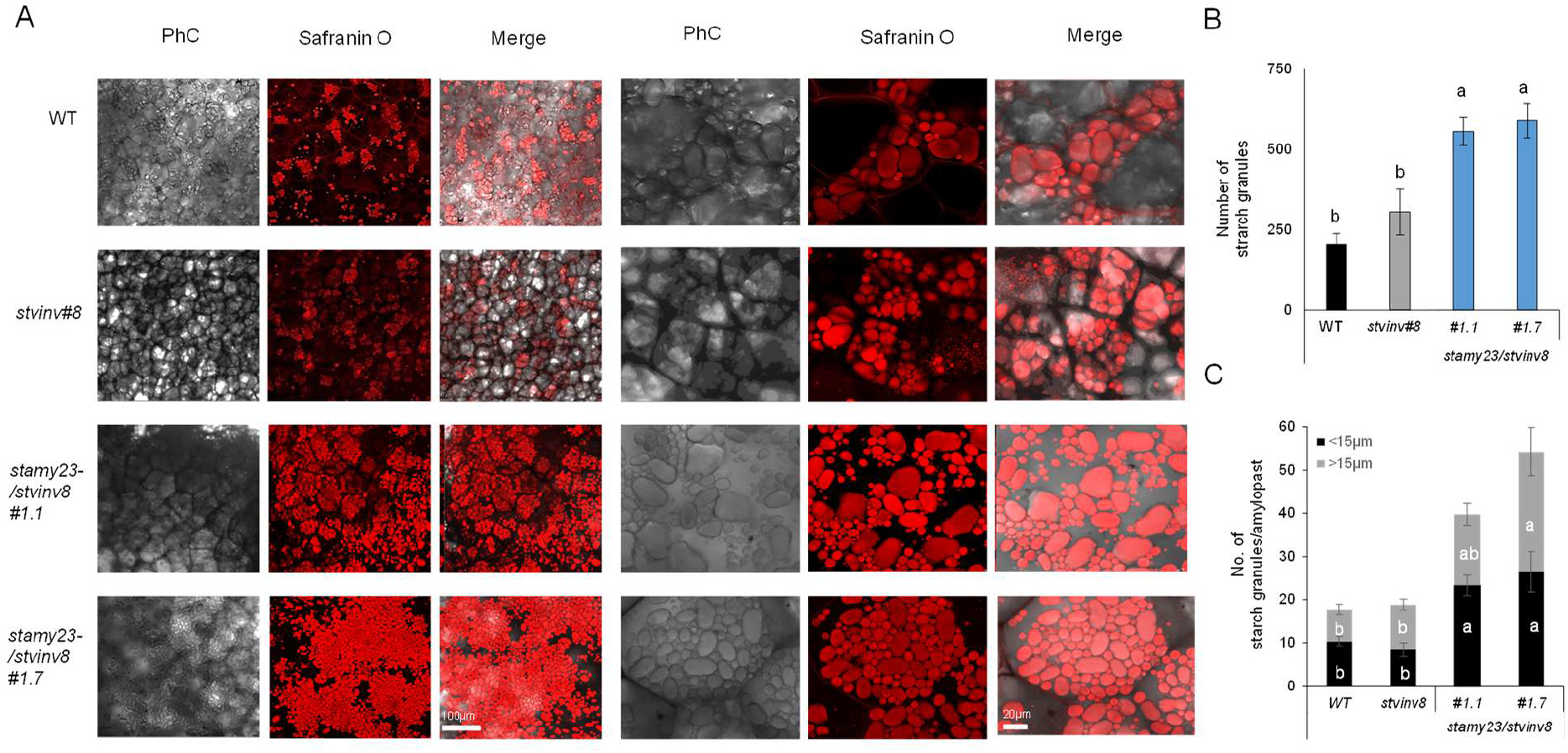
Starch-granules excess phenotype of *stamy23/stvinv#8* double mutant as compared to wild type (WT). (A) Starch granules in amyloplasts as observed with a confocal laser microscope 14 days after harvest. The images compare WT and single mutant (*stvinv#8*) to *stamy23*/*stvinv8#1*.*1, 1*.*7* double-knockout lines. Phase contrast (PhC), Safranin O staining, and merged images are presented. The scale bar equals 100µm and 20µm in the left and right panels, respectively. (B-C) Quantification of starch granules in a square of 500µm^2^ (B), and the number of starch granules per amyloplast categorized by size (C) analyzed from figure A, using Imaris software. Error bars represent standard error (SE). Different letters indicate significant differences between treatments at each time point (one-way ANOVA, p < 0.05). ANOVA, analysis of variance.

## Discussion

### *StAMY23* knockout delays dormancy release

Among the major challenges in potato storage are CIS and premature sprouting associated with dormancy release. We previously proposed that ED duration correlates with the accumulation of soluble sugars that act first as a signal and later as an energy source supporting bud outgrowth [12]. Using CRISPR/Cas9-mediated mutagenesis, we successfully generated two independent knockout lines *stamy23#11* and *stamy23#13* (Fig. 1). Dormancy release analysis, following cold treatment, revealed that both mutant lines exhibited prolonged ED (Fig. 2), strongly supporting a role for *StAMY23* in dormancy regulation. Consistent with this model, loss of *StAMY23* function inhibits starch degradation, thereby limiting both the signaling function and metabolic resources required to initiate sprouting. Interestingly, despite its role in dormancy regulation, knockout of *StAMY23* (#11 and #13) did not affect the starch granule density and size within amyloplasts as compared to WT (Fig. 2). This observation aligns with previous findings that StAMY23 enzyme primarily targets soluble cytosolic glucans, such as phytoglycogen, rather than insoluble starch granules within plastids. Its cytosolic localization supports this proposed role for the StAMY23 protein [20]. The broader significance of α-amylase in regulating dormancy and germination is well-established across plant species. In cereals, α-amylase is a key factor of pre-harvest sprouting, a process triggered when mature grains encounter excess moisture, breaking seed dormancy and causing premature germination [29]. Genetic studies further emphasize this relationship: introducing a barley α-amylase inhibitor into wheat delays germination. Transcriptome analyses show higher expression of α-amylase inhibitor genes in dormant wheat seeds compared to those that have released dormancy [30, 31]. Conversely, overexpression of *TaAMY1* in developing wheat grains reduces dormancy by promoting accumulation of soluble sugars such as α-gluco-oligosaccharides and sucrose [32]. Similar findings are reported in other species: suppression of *RAMY1A* in rice delays germination [33], and in potato, RNAi silencing of *StAMY23* extends tuber dormancy and delays sprouting by one to two weeks compared to WT [21]. Collectively, these studies underscore a conserved and central role for α-amylase in controlling the transition from dormancy to active growth.

### *stamy23* knockout in a *stvinv* background leads to induce starch accumulation

Tubers of the *stamy23/stvinv#8* and *stamy23/stvinv#21* double mutants exhibited similar ED and CIS tolerance, comparable to the corresponding single *stvinv#8* and *stvinv#21* mutants (Figs. 5 and 6). However, the double mutants accumulated higher starch levels than both the single mutants and WT plants (Fig. 7; Fig. S1), indicating that increased starch retention requires the combined loss of StAMY23 and StVINV enzymes activity.

In potato tubers, starch accumulates during development and serves as the main carbon reserve supporting sprout growth after dormancy release [34]. Sucrose imported from source leaves is converted in tuber cells to glucose 6-phosphate (G6P), which is transported into amyloplasts and converted to ADP-glucose, the direct precursor for starch biosynthesis [3]. Thus, final starch content reflects the balance between sucrose utilization, starch degradation, and carbon consumption during storage and sprouting. Downregulation of *StVINV* elevates sucrose and reduces hexose accumulation during postharvest cold storage, a metabolic state associated with reduced CIS and altered ED release. When temperatures permit bud growth, accumulated soluble sugars can be rapidly consumed to support respiration and sprouting [12, 16, 34]. In parallel, knockout of *StAMY23*, which is involved in degradation of soluble cytosolic glucans such as phytoglycogen and malto-oligosaccharides, may further limit carbon mobilization [35]. Therefore, the combined *stamy23/stvinv* mutation appears to shift tuber carbon metabolism toward starch conservation rather than sugar consumption, maintaining a storage-oriented metabolic state [36]. These findings extend previous potato starch-engineering studies, which mainly focused on modifying starch composition rather than increasing total starch accumulation. Editing of *GBSSI, SBEI, SBEII, SSII*, and *SSIII* altered amylose content, branching structure, and starch physicochemical properties, but did not substantially enhance starch yield [37, 38]. These results support the view that single-gene modification often fails to enhance total starch accumulation because starch biosynthesis is controlled by multiple genetic and enzymatic factors [39]. By contrast, tuber-specific overexpression of sucrose synthase increased ADP-glucose, starch accumulation, and tuber yield, highlighting the importance of sucrose-to-starch conversion in determining sink strength [40]. Together, our results show that the combined *stamy23/stvinv* mutations increases starch accumulation while improving storage performance and reducing CIS. This suggests that targeting complementary pathways controlling sucrose utilization and glucan degradation is more effective than editing single starch biosynthetic or structural genes, extending starch engineering from modifying starch composition toward enhancing starch retention and storage stability.

### Selected *stvinv* backgrounds may serve as platforms for improving agrotechnical potato properties

Targeted CRISPR/Cas9 mutagenesis of *StVINV* in elite potato cultivars showed that even partial allele knockout substantially reduced CIS, producing significantly lighter chip fry color, a key trait for the processing industry (Fig. 3 and 6). The mutant lines also exhibited prolonged ED, reducing sprouting-related losses and extending the market window for fresh tubers (Fig. 3 and 5). Importantly, these improvements were not accompanied by major effects on plant growth, canopy architecture, or tuber formation, indicating that *StVINV* modification does not impose a clear agronomic penalty.

Beyond storage and processing traits, *stvinv* lines showed improved cold and drought tolerance, likely associated with elevated sucrose accumulation. Sucrose may contribute both as a protectant and as a signaling molecule regulating growth, development, hormone pathways, and stress response [25, 26]. Sugars not only serve metabolic functions but also act as central signaling molecules that regulate plant growth, development, and responses to biotic and abiotic stresses. Variations in sugar type, concentration, and cellular energy status influence cell division, developmental programs, and stress adaptation pathways. Consistent with this regulatory role, sucrose has been linked to stolon and tuber development in potato through auxin, cytokinin, and gibberellin signaling, and to bud outgrowth and branching in potato and other species. Together, these findings highlight *StVINV* as a central regulator of tuber physiology, storage behavior, and stress adaptation [41]. The combination of *stamy23* and *stvinv* further demonstrates that targeting complementary metabolic pathways can produce stronger effects than single-gene modification. Coordinated suppression of sucrose utilization and glucan degradation enhanced starch accumulation while maintaining improved storage traits and reduced CIS. This supports the broader principle that multigene engineering is often required to improve complex agronomic traits controlled by interconnected metabolic and regulatory networks [42].

## Material and Methods

### Plant material, transformation and sample collection

The plant material of *S. tuberosum* L. cv. Désirée (2n = 4x = 48) used in this study was propagated in vitro every 6–8 weeks in 1× Murashige and Skoog (MS) medium (pH 5.8) containing vitamins (Duchefa, Haarlem, the Netherlands), 3% (w/v) sucrose, 0.8% (w/v) Phyto agar (Duchefa), and 4μM silver thiosulfate. Plants were cultivated in a growth chamber at 25°C with a 16h light/ 8h dark photoperiod.

Potato leaves were used for Agrobacterium-mediated leaf disk infection [43]. Transgenic plants were screened on 50 mg/l kanamycin or 12 mg/l hygromycin (Duchefa). Protoplasts were isolated from 1 gr of leaves of 4-week-old cv. Désirée plants, transfected, and regenerated into shoots as previously described [25]. Seedlings were propagated in tissue culture for 3 weeks and then transferred to small pots (4.5 × 8.5 cm) containing standard potting mix (Pelemix Green) for an additional 6 weeks. Seedlings were maintained in a controlled growth chamber under a 16 h light/8 h dark photoperiod at 23 °C, with ∼200 µmol m^−2^ s^−1^ photosynthetically active radiation provided by LED illumination. Subsequently, plants were transferred to the greenhouse under natural sunlight during short days. Watering was stopped 2 weeks before tuber harvest. After harvest, the tubers were incubated for 14 days at 14°C and 95% relative humidity to facilitate tuber skin curing. Then they were transferred to the specified temperature treatments. Samples were taken from the tuber parenchyma under the apical bud using a cork borer (diameter 1 cm, 3 cm penetration), immediately frozen in liquid N2, and transferred to −80°C. Samples of 1g were taken for sugar analysis.

### CRISPR/Cas9 sgRNA design and vector construction

Single guides RNA (sgRNA) were designed for *StVINV* (PGSC0003DMG400013856) and *StAMY23* (PGSC0003DMG400009891) using the CRISPR design tool http://crispr.hzau.edu.cn/CRISPR2/. For *StVINV*, sgRNA9: 5′-GGACACATCATATAACGGCC-3′ targeting exon 2 upstream of the NGG protospacer adjacent motif (PAM) was selected. It was cloned into the non-binary plasmid pSAT-Cas9-sgRNA(9) (8 kb), as previously described [25]. For *StAMY23*, two sgRNAs were designed: sgRNA1: 5′-GGCCACACAGTCAGTAACCG-3′ and sgRNA2: 5′-GTTGGGACTACTCAAGGGCA-3′ targeting exon 3 downstream of a GGA PAM and exon 4 upstream of a TGG PAM, respectively. The sgRNA expression cassettes were synthesized into pUC57-KANA. The CRISPR components were then assembled using the GoldenBraid modular cloning system [44]. The final binary constructs included a plant codon-optimized Cas9 driven by the StUBIQUITIN10 promoter (proStUBIQ) as previously described [45].

### Mutant screening and genotyping

Genomic DNA was isolated from T0 potato plants according to Dellaporta et al. [46]. Mutant screening was performed using primers flanking the target regions: VInvF 5′-ACCATCCTACCCGATGGTCA-3′ / VInvR 5′-CAGGTCAGCAGATTCACTAT-3′ and AMY23F 5′-AGATTACGAGCTCTAGCAAT-3′ / AMY23R 5′-TGGCGAATAGAAGTAGGTGG-3′. PCR products were digested with *BSURI* and *MspI*, respectively, and separated on a 1% agarose gel. Shifts in band size or undigested products compared to WT indicated potential mutations.

Genotyping was confirmed via Amplicon-seq NGS (Syntezza, Jerusalem) or PLASMIDSAURUS premium PCR sequencing (https://plasmidsaurus.com/premium_PCR_sequencing).

### Dormancy duration

Endodormancy duration was measured by storing the tubers at 14°C for 14 days to cure harvest wounds. This was followed by storage at 4°C for two or four weeks to enhance ED release and then incubation at 14°C until sprouting. Dormancy was considered released when the apical bud exceeded 2 mm in length.

### Sugar extraction and quantification

Sucrose, glucose, and fructose levels were extracted and quantified as previously described [25]. Briefly, bud-base parenchyma tissue was sampled using a cork borer (Ø 1 cm, 3 cm penetration), weighed, and immediately frozen in liquid N2 and transferred to −80°C until use. Tissues were incubated three times in 80% ethanol at 80°C for 45 min each time. The solution was dried using a speed vacuum (Centrivap concentrator, Labconco) and passed through a 0.2-mm membrane filter (Millex-GV filter unit, Merck Millipore). The filtrate was used for sucrose, glucose, and fructose analyses by ultrafast liquid chromatography (UFLC) in an LC-10A UFLC series system (Shimadzu) equipped with a SIL-HT automatic sample injector, pump system, refractive index detector (SPD-20A), differential refractometer detector (Waters 410) and analytical ion-exchange column (6.5 × 300 mm, Sugar-Pak I, Waters). The mobile phase (ultrapurified deionized water, Bio-Lab) was eluted through the system for 20 min at a flow rate of 0.5 mL/min, and the column temperature was set to 80°C. The chromatographic peak corresponding to each sugar was identified by comparing the retention time with that of a standard. A calibration curve was prepared using standards to determine the correlation between peak area and concentration.

### Starch staining and extraction

Safranin-O staining was performed as described by Liu et al. [47]. Granules counts were done using laser confocal scanning microscopy and Bitplane Imaris software version 8.0.1 (Bitplane A.G.). About 100 cells in three biological replicates were analyzed per genotype. Starch extraction was performed using 2M HCl hydrolysis for 2 hours at 100°C [48]. After neutralizing the solution to pH 7 with NaOH, a Sumner reaction was conducted [49]. Absorbance was measured at 550nm and compared against the glucose concentration standard curve. To express the experimental results in terms of starch concentration, the glucose concentration values were multiplied by 0.9.

---

**Figure S1.**
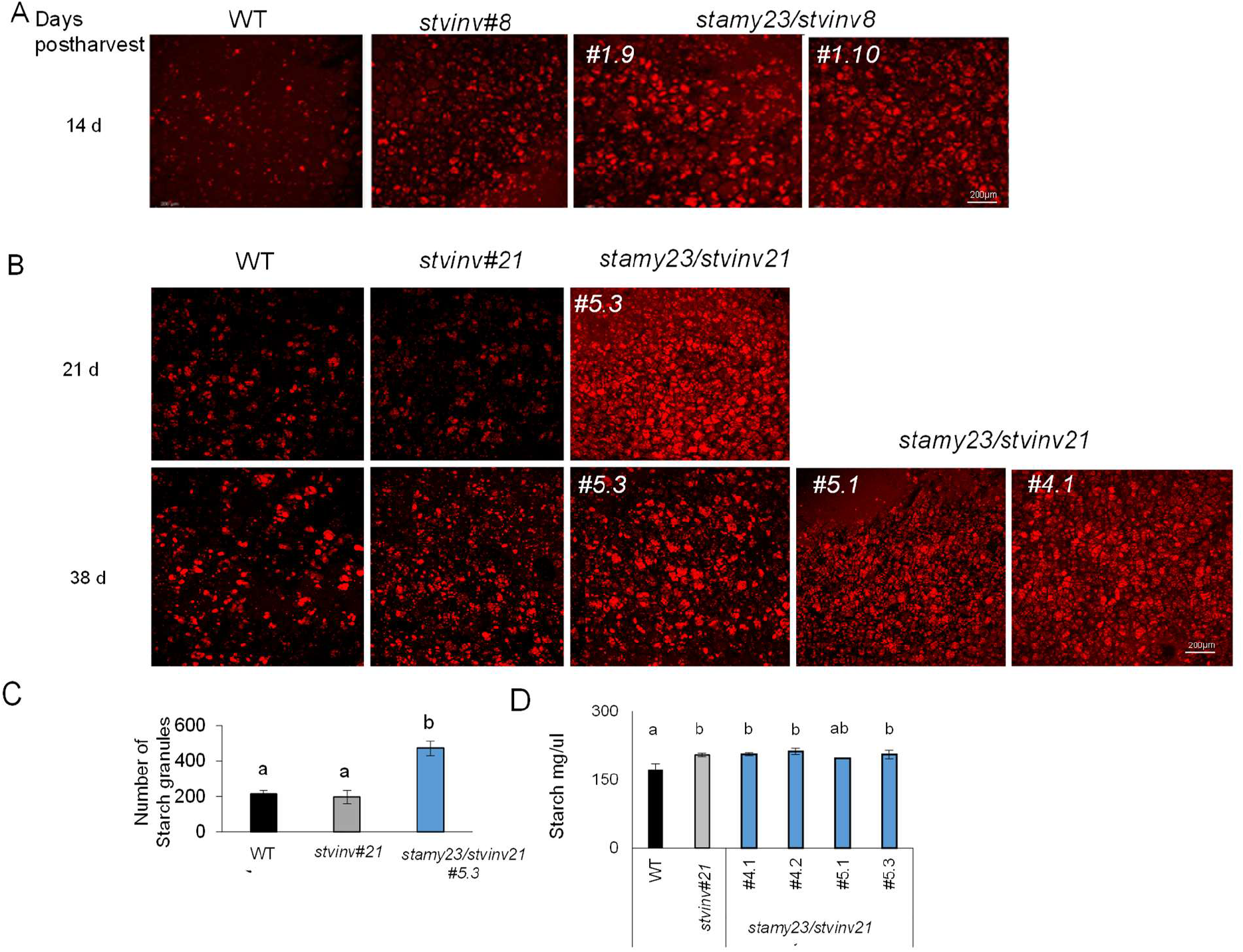
Starch granules excess phenotype of *stamy23/stvinv#8* and *stvinv#21* double mutants. (A-B) Starch granules within amyloplasts, stained with Safranin O, were observed using a confocal laser microscope. (A) The images compare wild type (WT) and single mutants (*stvinv#8*) to *stamy23/stvinv8 #1*.*9 and #1*.*10*, 14 days postharvest and (B) *stvinv#21 5*.*3# to stamy23/stvinv21#4*.*1, #5*.*1, #5*.*3* double-knockout lines, 21, and 38 days postharvest. The scale bar equals 200µm. (C) Quantification of starch granules from (B; upper panel) and (D) from (B; lower panel) using Imaris software. Error bar represents standard error (SE). Different letters indicate significant differences between treatments at each time point (one-way ANOVA, p < 0.05).

## Notes

### Competing Interest Statement

The authors have declared no competing interest.

